# Effect of Methamphetamine on Ultraweak Photon Emission and Level of Reactive Oxygen Species in Male Rat Brain

**DOI:** 10.1101/2022.09.14.508017

**Authors:** Tahereh Esmaeilpour, Azam Lotfealian, Morteza Anvari, Mohammadreza Namavar, Narges Karbalaei, Abbas Shahedi, Istvan Bokkon, Hadi Zadeh-Haghighi, Christoph Simon, Vahid Salari, Daniel Oblak

**Author notes:** These authors contributed equally to this work (Tahereh Esmaeilpour, Azam Lotfealian).

## Abstract

All living cells, including neurons, generate ultra-weak photon emission (UPE) during biological activity, and in particular, in the brain, it has been shown that UPE is correlated with neuronal activity and associated metabolic processes. Various intracellular factors, as well as external factors, can reduce or increase the intensity of UPE. In this study, we have used Methamphetamine (METH) as one potentially effective external factor, which is a substance that has the property of stimulating the central nervous system. METH can impair mitochondrial function by causing toxicity via various pathways, including an increase in the number of mitochondria, hyperthermia, the increased metabolic activity of the brain, and the production of glutamate and excess calcium. In addition to mitochondrial dysfunction, METH alters cellular homeostasis, leading to cell damage and the production of excess ROS. The aim of this study is to measure and compare the UPE intensity and reactive oxygen species (ROS) levels of the prefrontal, motor, and visual cortex before and after METH administration. Twenty male rats were randomly assigned to two groups, the control, and METH groups. In the control group, 2 hours after injection of normal saline and without any intervention, and in the experimental group 2 hours after IP injection of 20 mg/kg METH, sections were prepared from three areas: prefrontal, motor, and V1-V2 cortex, which were used to evaluate the emission of UPE using a photomultiplier tube (PMT) device and to evaluate the amount of ROS. The results showed that the amount of ROS and UPE in the experimental group in all three areas significantly increased compared to the control group. So, METH increases UPE and ROS in the prefrontal, motor, and visual regions, and there is a direct relationship between UPE intensity and ROS production. Therefore, UPE can be used as a dynamic reading tool to monitor oxidative metabolism in physiological processes related to ROS. Also, the results of this experiment can support the hypothesis that the production of excess UPE may lead to the phenomenon of phosphene and visual hallucinations.

## 1 Introduction

### 1.1 Ultraweak Photon Emission (UPE)

It has been shown that almost all living systems spontaneously emits photons with a very low intensity with a flux of about a few to thousands of photons/cm^2^ per second at wavelengths mostly in the range of 200-800 nm, called ultraweak photon emission (UPE) [1–3], which is an entirely distinct phenomenon from other known biological light emission processes, for instance, it is different from black body radiation (i.e. infra-red thermal energy) and bioluminescence (e.g. from Luciferin-Luciferase reactions or photoprotein processes) [4]. UPE is in fact expressed in other living tissue/organs (e.g. hand, face) and other living systems (e.g. plants, bacteria), however, specifically in the brain, UPE is correlated with neuronal activity and associated metabolic processes, e.g. it correlates with EEG activity, cerebral blood flow, hyperoxia, glutamate release, neuronal-glia calcium homeostasis, as well as, metabolism of the inner mitochondrial respiratory chain through the production of reactive oxygen species (ROS) [5–8], and the origin of UPE is in direct connection with ROS [1, 2]. The UPE intensity variations are associated with different physiological and pathological conditions, e.g. thermal, chemical and mechanical stress, mitochondrial respiratory chain, cell cycle, etc.

### 1.2 Methamphetamine

Methamphetamine (METH; N-methyl-alpha-methylphenethylamine) is a strongly addictive synthetic psychostimulant that is widely abused in the world. METH has high lipophilicity, which makes it possible to easily cross neuronal membranes and the blood-brain barrier (BBB). In addition, the chemical structure of METH is similar to that of monoamines. As a result of this, dopamine (DA), serotonin (5-HT), and the nor-adrenaline plasma membrane transporter can transport METH into neurons and neuronal terminals in the brain [9]. METH acts on the monoamine storage vesicles and depletes them of neurotransmitters. Namely, METH redistributes monoamines from storage vesicles into the cytosol [10]. In other words, in the central nervous system (CNS), METH stimulates the release and partially inhibits the reuptake of newly synthesized catecholamines. This mentioned process results in the release of DA, noradrenaline, and 5-HT into the synaptic cleft, which then stimulates postsynaptic monoamine receptors [11]. Although the exact mechanisms are not clear, studies suggest that disturbed mitochondrial mechanisms have a key role in methamphetamine neuropathology [12–15]. It is probable that the METH-induced high level of cytoplasmic DA produces an unregulated overproduction of ROS and oxidative stress on the neuron [16]. In addition, METH inhibits monoamine metabolism via complex inhibition of mitochondrial monoamine oxidase (MAO) [17]. METH perturbs the mitochondrial membrane potential and electron transport chain, so METH can kill neurons not only by the direct generation of ROS but also by initiating the mitochondrial-dependent induction of apoptotic cascades [15]. Studies also revealed that chronic METH abuse can produce neurodegenerative changes in the human brain, such as damage to dopamine and serotonin axons, loss of gray matter, and microgliosis in various brain regions [18]. METH consumption correlates with reduced neuronal integrity and viability and active neurodegeneration, mainly in dopamine pathways, which can cause serious mental illness [19]. METH-induced serious mental illness like psychosis includes delusions; auditory, visual, and tactile hallucinations; depressive syndrome; paranoia, a tendency toward violence; and an increased risk of suicide, among others [20–24].

### 1.3 Reactive Oxygen Species (ROS)

Reactive oxygen species (ROS) are generated naturally as a consequence of cellular aerobic metabolism in a biological system. Mitochondria, plasma membranes, endoplasmic reticulum, and peroxisomes are the principal sites of ROS generation within the cell and mitochondrial respiration is a major source of ROS. On the other hand, Oxidative stress (OS) happens when ROS are created in excess or when cellular defences are unable to metabolize them [25]. Because of the existence of polyunsaturated fatty acids (PUFAs), cell membranes are vulnerable to oxidative damage. During oxidative stress, ROS attack lipids with carbon-carbon double bonds, particularly PUFAs, and this causes membrane lipid peroxidation which leads to malondialdehyde (MDA) production. Consequently, MDA is one of the end products of polyunsaturated fatty acid peroxidation in cells and a biomarker for lipid peroxidation, and an increase in free radicals leads to an increase in MDA production [26, 27]. Hydroxyl radical (HO•) and hydroperoxyl (HOO•) are the two most common ROS that can have a significant impact on lipids. The hydroxyl radical (HO•) is a chemically reactive species of activated oxygen which can be formed from O_2_ during cell metabolism and under various stress situations. the hydroperoxyl radical (HO•_2_) is the protonated form of superoxide and plays a crucial role in lipid peroxidation. This produces H_2_O_2_ which can react with redox active metals such as iron or copper to produce the hydroxyl radical (HO•) via Fenton or Haber-Weiss reactions [28].

In this research, we measure and compare the UPE intensity and ROS levels of the prefrontal, motor, and visual cortex before and after METH administration. The variations in the UPE intensity may have a potential applications as indicators for cognitive impairment studies and medical diagnostics.

### 2 Materials and Methods

### 2.1 Samples

20 animals were allocated into two groups (n=10 per group). 2 hours after injection of normal saline in the control group and methamphetamine with an optimized dosage 20 mg/kg in the experimental group intraperitoneally, the animals were sacrificed under deep anesthesia with ether and their brain was removed and placed on ice and in a solution of artificial cerebrospinal fluid (ACSF). The duration of anesthesia was short and the application of low dose ether was similar in all cases. So, the effect of ether is almost excluded, and therefore the effect of METH is discriminated with the control groups. Sections of three areas V1-V2, prefrontal, and Motor with a thickness of 0.5 mm were prepared by Brain Matrix. The slices are divided into two groups, for evaluating the UPE using the PMT device and the amount of ROS using the molecular index of malondialdehyde (MDA) by the TBARSassay method and calorimeter by the ELIZA Reader device. The prepared ACSF was refrigerated to cool during use and 95% O_2_ and 5% CO_2_ were passed through at a rate of 2.5 ml/min.

### 2.2 Photon counting

A photomultiplier tube (PMT) is an extremely sensitive detector that can detect and count single photons. A PMT system (Hamamatsu Photonics K.K., Electron Tube Center, Hamamatsu, Japan, R6095 PMT providing a maximum spectral response from 300 to 700 nm with a maximum detection at 420 nm with about 30% quantum efficiency) has been used in our experiments to observe time-dependent photon emission intensity in the dark box located in a dark room. The distance between the sample and the PMT was approximately 1 cm. The PMT is connected to a counter, which is also connected to the computer for data to be digitally visible. The specimen will be located in the dark room and the electronic equipments will be placed outside the dark room, so that no other light except the sample UPE could be measured by PMT (see Figure 2) [3].

**Fig 1.**
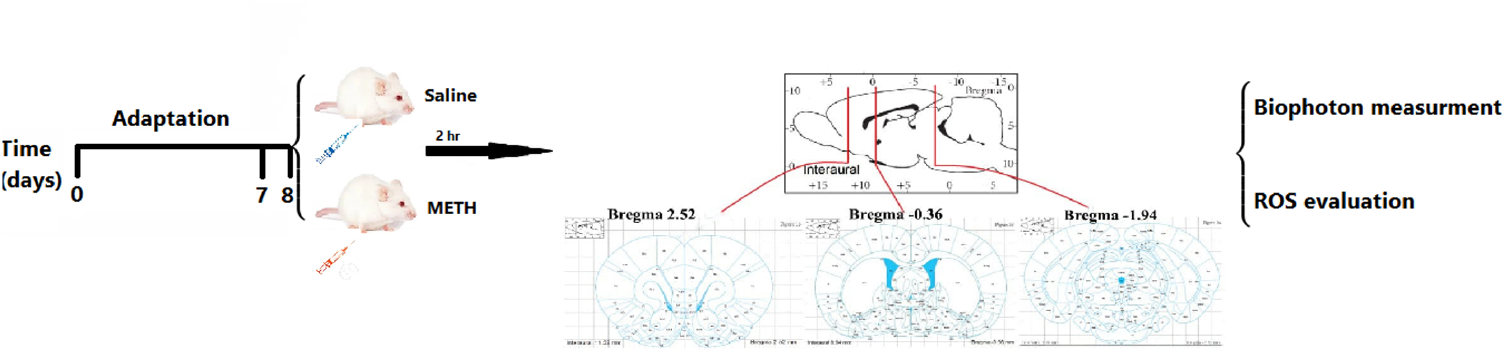
Schematic diagram of the experimental design. Two groups of animals have been studied. The control group, 2 hours after the injection of normal saline and without any intervention, and the experimental group 2 hours after the IP injection of 20 mg/kg METH. Sections were prepared from three areas of the brain: prefrontal, motor, and V1-V2 cortex, which were used to evaluate the emission of UPE using PMT device and to evaluate the amount of ROS.

**Fig 2.**
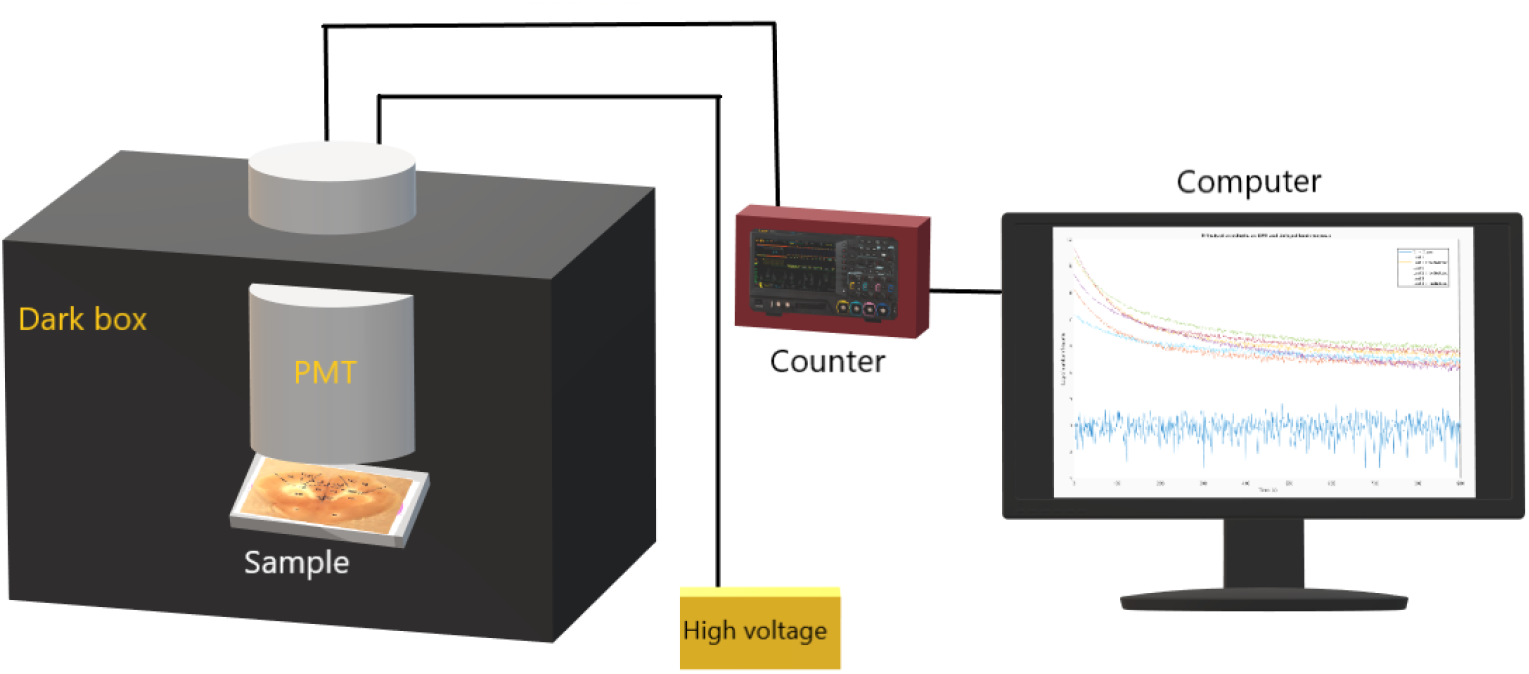
Ultraweak photon emission (UPE) from brain slices is measured in a dark box with a photomultiplier tube (PMT) which is a single photon detector.

### 2.3 Measurement of ROS level

Lipids are very sensitive to ROS attack and there are some biomarkers that can assess the amount of ROS production in cells, such as malondialdehyde (MDA), 4-hydroxy-2-nonenal (4-HNE) and F2-isoprostane, which are known biomarkers for lipid peroxidation assessment [30]. The measurement of ‘Thiobarbituric Acid Reactive Substances’ (TBARS) is a well-established method for monitoring lipid peroxidation. The TBARS assay detects the level of MDA, the major lipid oxidation product (see Fig.3), which is often considered a good index of the level of oxidative stress in a biological sample. The assay involves the reaction of lipid peroxidation products, primarily MDA, with thiobarbituric acid (TBA), which leads to the formation of MDA-TBA2 adducts called TBARS. TBARS yields a red-pink color that can be measured spectrophotometrically at 532 nm. The TBARS assay is performed under acidic conditions (pH = 4) and at 95 °C. Pure MDA is unstable, but these conditions allow the release of MDA from MDA bis(dimethyl acetal), which is used as the analytical standard in this method. The TBARS assay is a straightforward method that can be completed in about 2 h [29].

**Fig 3.**
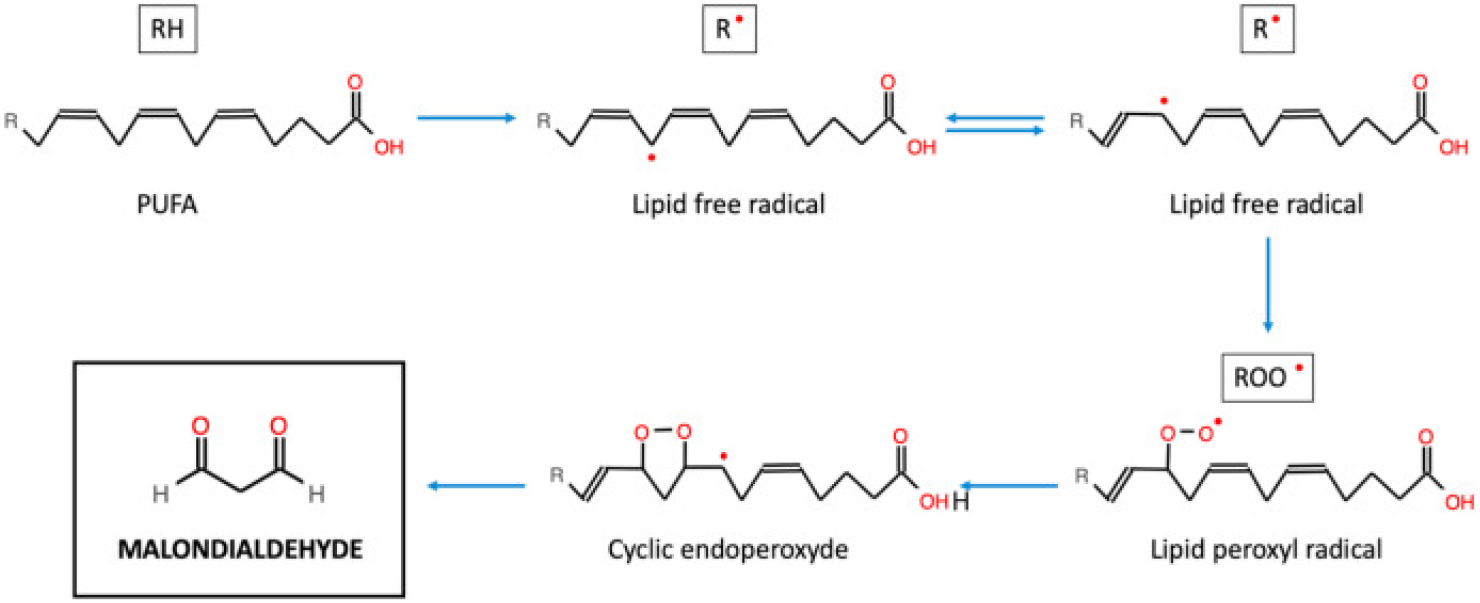
Malondialdehyde (MDA) formation through lipid peroxidation in mitochondria. Lipids are very sensitive to ROS attack and MDA is a biomarker for lipid peroxidation assessment. The figure is taken from the Ref. [30].

#### 2.3.1 Statistics analysis

Data analysis was performed using one-way ANOVA (post hoc: Tukey) and Unpaired t-test as appropriate (Prism 6; Graph pad Software Inc., San Diego, CA). Results were expressed as mean *±* SEM and the level of significance was set at a P value of less than 0.05.

#### 2.3.2 Ethical approval

All procedures performed in studies involving animals were in accordance with the ethical standards of the Ethics Committee (ir.ssu. medicine rec.1400.317), Yazd University of Medical Sciences.

## 3 Results

### 3.0.1 Optimization of methamphetamine concentration

Doses of methamphetamine were optimized by injecting 0.5 cc of 6 different concentrations of methamphetamine containing 15, 18, 20, 23, 25, 30, mg/kg on consecutive days for one week, dose 20 was the highest dose in which rats all survived and were most active.

Figure 4 shows the Photomicrographs of three representative coronal sections of the prefrontal, motor and visual cortex of the rat brain and anatomical labels according to the Paxinos and Watson atlas.

**Fig 4.**
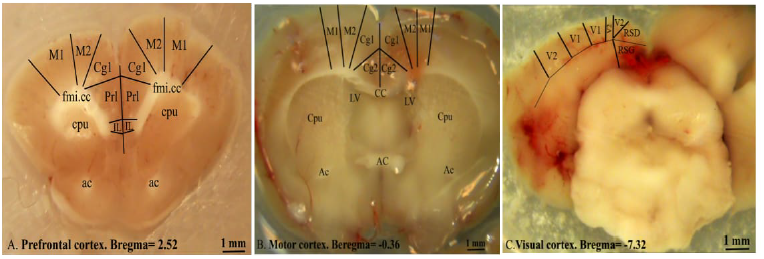
Coronal section of the A: prefrontal (Bregma 5.16 to 1.56 mm), B: motor area (M1, M2) (Bregma 5.16 to 3.24 mm) and C: visual area (V1-V2) (Bregma 4.20 to-9.36 mm). M1: primary motor/ M2: secondary motor/ Cg: cingulate cortex/ Prl: prelimbic cortex/ Il: infralimbic cortex/ Dp: dorsal peduncular cortex/ Cpu: caudate putamen/ ac: anterior commissure/ fmi: forceps minor of the corpus callosum / Cc: corpus callosum/ Lv: lateral ventricle/ V2: secondary visual cortex / V1: primary visual cortex.

### 3.0.2 Measurement of MDA in control and experimental groups

The results showed that the MDA level was significantly higher in the prefrontal and motor cortex compared to the visual cortex in the control group (Fig. 5A), but in the experimental group after METH administration only motor cortex showed that the MDA level was significantly higher relative to visual cortex (Fig. 5B).

**Fig 5.**
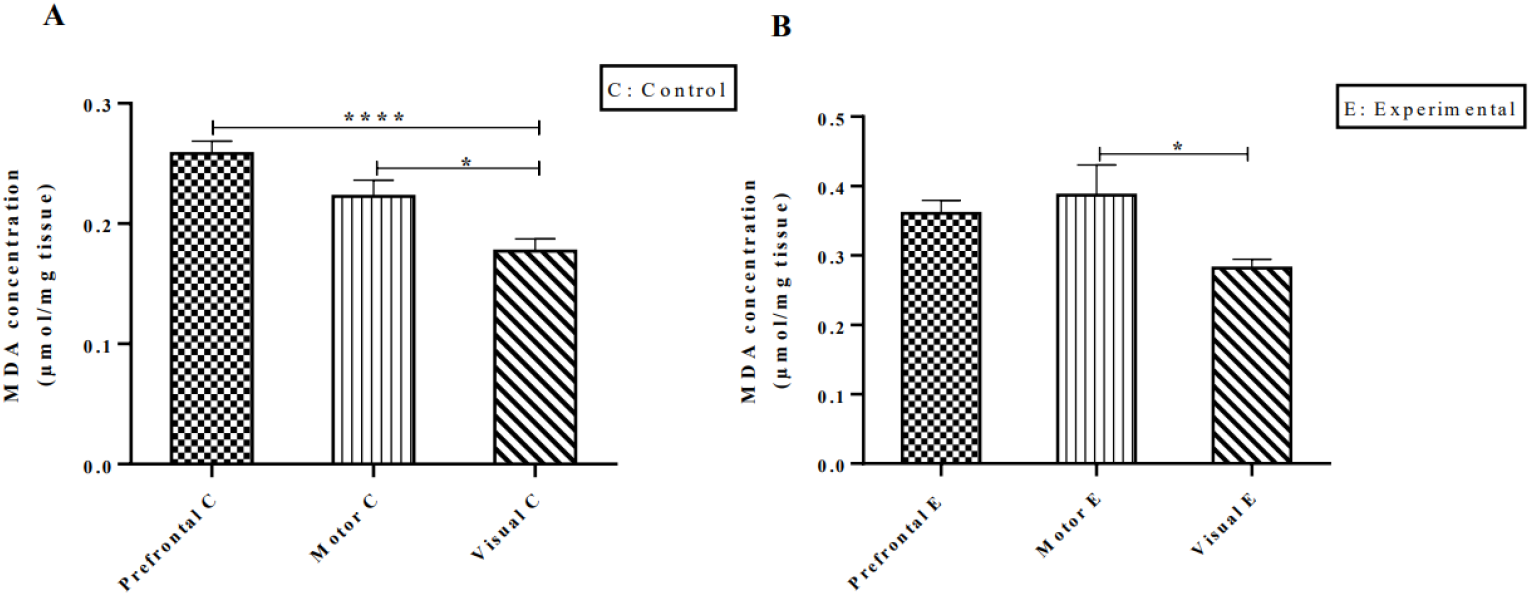
MDA production in prefrontal, motor and visual cortices in the control group (A), and methamphetamine (METH) group (B) (Mean ± SEM; n = 10 independent experiments; *p*<*0.05; ****p*<*0.0001; one-way ANOVA).

### 3.0.3 Comparison of MDA measurements in control and experimental groups

The results showed that the MDA production in the experimental group in all three areas was significantly increased compared to the control group. (Fig. 6)

**Fig 6.**
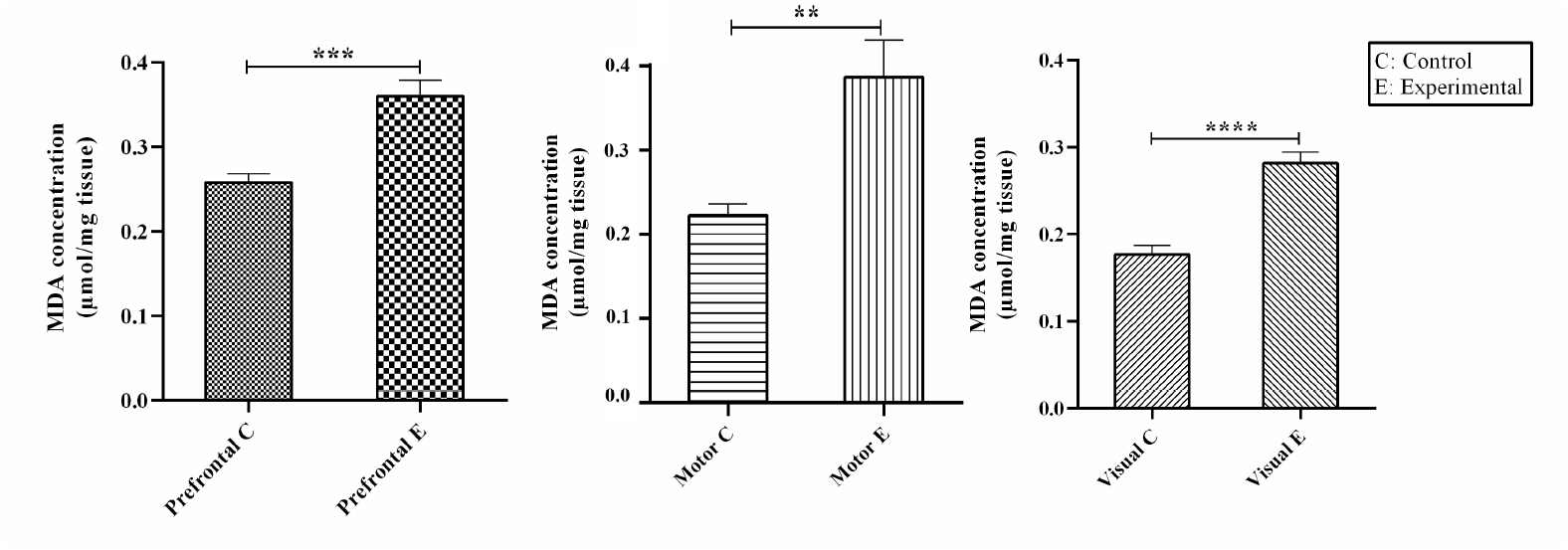
MDA alterations in prefrontal, motor and visual cortices in control and experimental groups. (Mean ± SEM; n = 10 independent experiments; **p*<*0.01; ***p*<*0.001; ****p*<*0.0001; Unpaired t-test; MDA= Malondialdehyde).

### 3.0.4 UPE measurements in control and experimental groups

The UPE in the control group did not show any significant difference between the three areas (Fig. 7A), but after methamphetamine administration, the motor cortex showed a significant increase compared to visual and prefrontal cortex (Fig. 7B).

**Fig 7.**
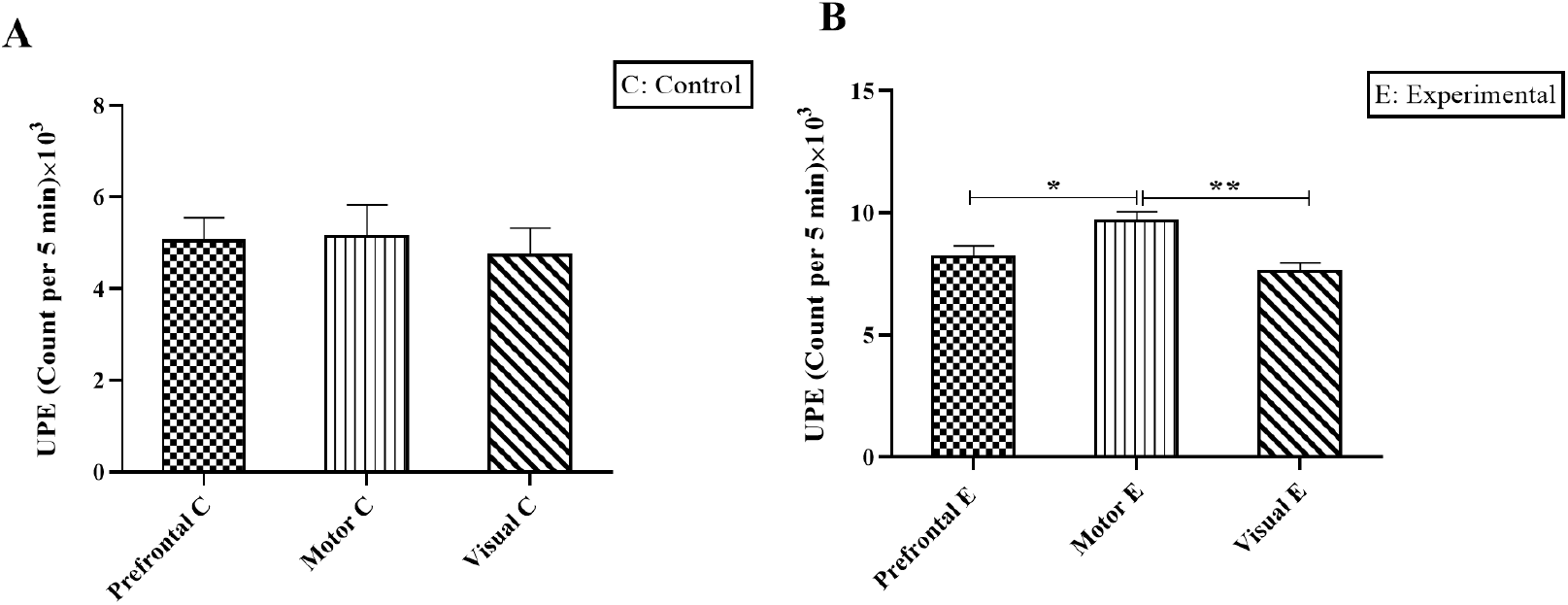
UPE intensity in the control group (A) and the experimental group (B) in prefrontal, motor, and visual cortex. (Mean ± SEM; n = 10 independent experiments; *p*<* 0.05; ** p*<* 0.01; UPE= Ultra-weak Photon Emission; one-way ANOVA).

### 3.0.5 Comparison of UPE levels of control and experimental groups

Methamphetamine also significantly increased the UPE intensity in all three groups. The enhancement in UPE intensity for motor cortex is more significant (Fig. 8).

**Fig 8.**
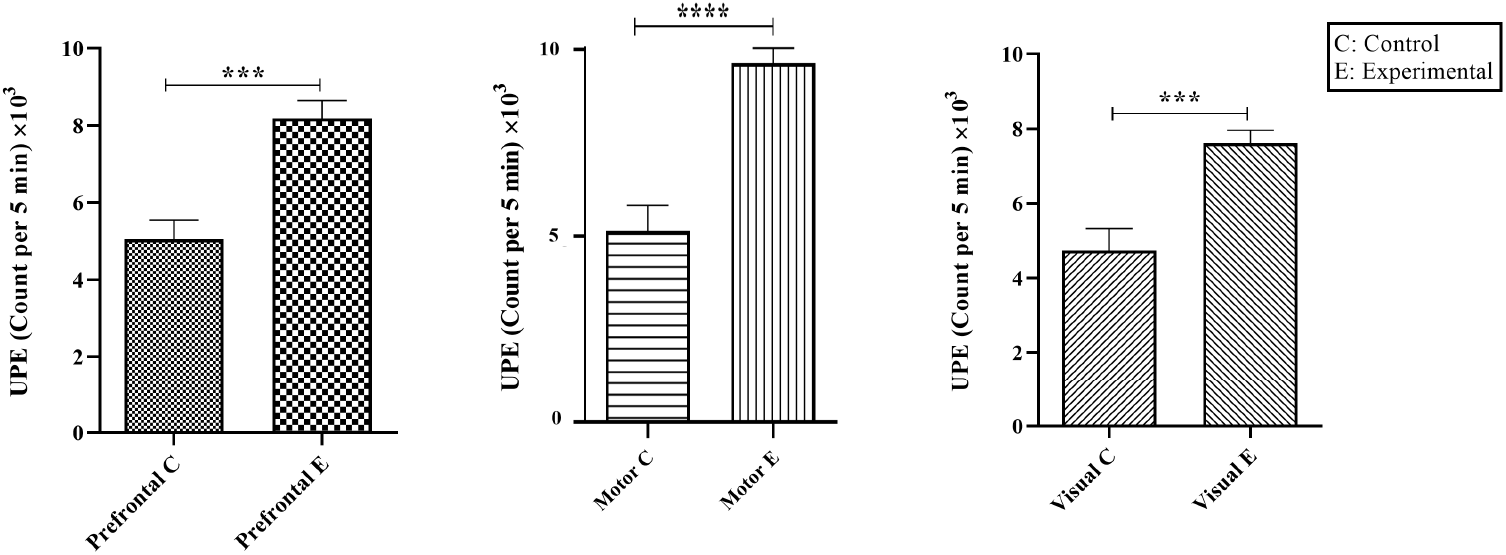
UPE alterations caused by methamphetamine administration in the experimental groups in prefrontal, motor, and visual cortex, increased significantly compared to the control groups. (Mean ± SEM; n = 10 independent experiments; *** p*<* 0.001; **** p*<*0.0001; Unpaired t test). UPE= Ultra-weak Photon Emission).

## 4 Discussion

In this study, methamphetamine injection showed a significant increase in the amount of ROS and UPE in the brain tissue of the experimental groups compared to the control group. The most important source of ROS production is the electron transfer chain in mitochondria [31]. Methamphetamine can impair mitochondrial function by causing toxicity from various pathways, including an increase in the number of mitochondria [32], hyperthermia, increased metabolic activity of the brain [33], and the production of glutamate [34] and of excess calcium [35, 36]. In addition to mitochondrial dysfunction, methamphetamine alters cellular homeostasis, leading to cell damage and the production of excess ROS. And since many experiments have shown that ROS is an important source of UPE production [37], it can be inferred that methamphetamine consumption, by increasing ROS, produces excess UPE in the brain, which is observed in this study as well. In the results obtained in this study, the amount of ROS and UPE produced in the motor and prefrontal cortices was higher than in the visual cortex, which can be due to differences in the type of neurons, synaptic connections and their excitability threshold. The motor cortex, for example, contains specialized large pyramidal neurons. These large neurons are tightly connected and have more inputs and connections to other neurons [38]. Neurons with longer axons and multiple synapses have greater biological requirements for axonal transmission, leading to higher demand for ATP, which combined with partial mitochondrial dysfunction makes this group of neurons more vulnerable to oxidative damage [39]. On the other hand, in previous experiments by transcranial magnetic stimulation (TMS), it has been shown that the excitability threshold of the visual cortex is significantly higher than the motor cortex [40]. This difference can be due to the presence of more pyramidal cells in the motor cortex and the different effects of TMS on the function of neurotransmitters [41]. Also, by stimulation of the motor cortex and prefrontal by TMS, it has been shown that the electric fields caused by the stimulation of the motor cortex are higher than in prefrontal cortex and the excitability threshold of the prefrontal area is much higher than the motor area [42], maybe in line with the results of the present study, in which we showed that the effect of methamphetamine was greater on cells of the motor cortex that have higher excitability, which led to the production of more ROS and UPE in this area. The production of excess UPE in the retina and visual cortex may lead to phosphonic light (biophysical images), and these photons can be converted into biophysical images (phosphene) only in the V1 and V2 regions due to their retinotopic structure [43]. In addition, METH increases wakefulness and physical activity, therefore it may be possible that significant UPE from prefrontal and motor regions due to the METH induced increased cognitive instability and delusion (prefrontal region) and hyperactivity (motor region) [44]. The results of this study, in accordance with previous studies, shows that methamphetamine, as a psychoactive substance that causes visual hallucinations (e.g. phosphene), stimulates the nervous system, which in turn leads to an increase in ROS and thus the production of excess UPE. Our findings here are consistent with the hypothesis of a relationship between UPE and phosphene light production [43].

### 4.0.1 A hypothesis for the action of METH at the molecular and atomic level

Administration of METH is associated with behavioral hyperactivity and used to treat attention-deficit hyperactivity disorder (ADHD), which is accompanied by production of ROS [45]. METH belongs to the group of medicines called central nervous system (CNS) stimulants. On the other hand, Lithium studies have shown improvement in irritability, anger, and aggression in children with ADHD. Moreover, a recent study suggests that a radical pair model involving ROS and flavin can explain lithium effects on hyperactivity, where the lithium’s nuclear spin exerts its effects via modulating the spin dynamics of naturally occurring radical pairs in the brain [46]. It, thus, seems relevant to propose a quantum model based on radical pairs for the effects of methamphetamine on hyperactivity. It should be noted that nicotinamide adenine dinucleotide phosphate (NADPH)-oxidase (NOX) is a class of enzymes that is a major source of ROS production in the brain [47], and that the NOX enzymes are flavoproteins. The radical pair mechanism is a leading theory for how birds and other animals sense magnetic fields. In this context, flavin is a canonical component of the radical pairs [48]. Furthermore, it has been shown that exposure to transcranial magnetic stimulation modulates the behaviour of the methamphetamine-administered patients [49–53]. Magnetic field effects are abundant in biology, and it has been suggested that the radical pair mechanism may underlie many of these phenomena [54]. To explore radical pair related quantum effects, here, one can test the effects of applied magnetic field and isotope effects, e.g., using ^15^N in methamphetamine instead of the usual ^14^N. It should be noted that ^15^N and ^14^N carry nuclear spin of 1/2 and 1, respectively. (see Fig. 9).

**Fig 9.**
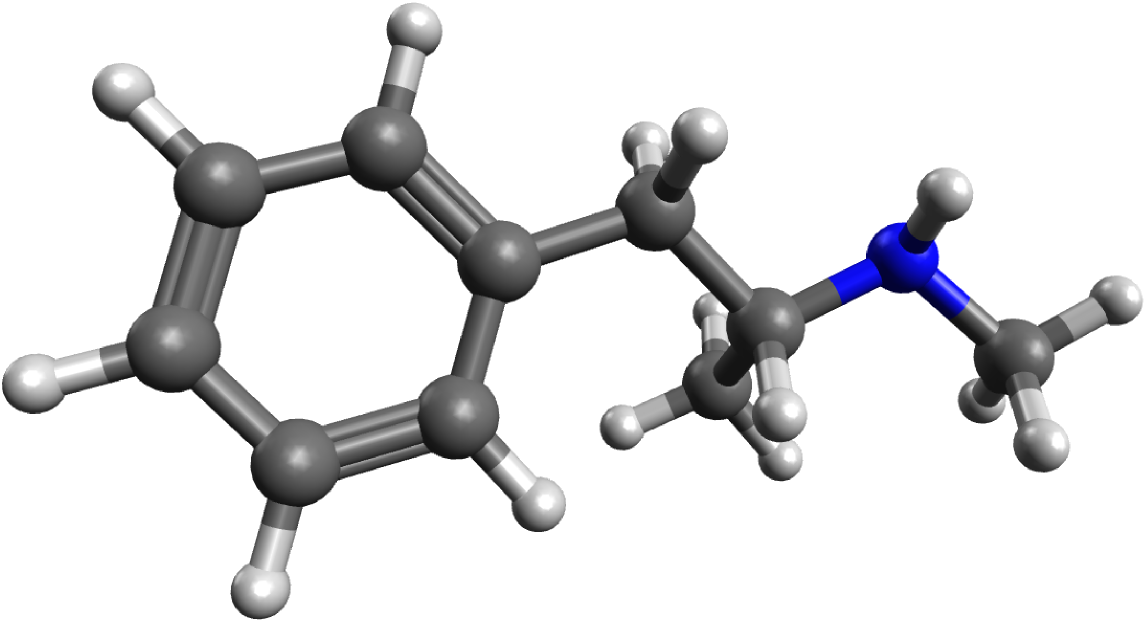
Molecular structure of methamphetamine (C_10_H_15_N). Blue ball is nitrogen.

## 5 Conclusion

In this study, we have found that methamphetamine injection in the experimental group significantly increases ROS and UPE compared to the control group. This increase was greater in the motor cortex, possibly due to differences in excitability thresholds, synaptic connections, light-sensitive chromophore molecules, the size and shape of neurons, the amount of neuronal activity induced by neurons, and the amount of ROS produced by mitochondria. Overall, it can be concluded that since ROS is the most important factor in UPE production, there is a strong correlation between UPE intensity and ROS production due to methamphetamine administration. This research confirms that UPE can be used as a dynamic reading tool to monitor oxidative metabolism in physiological processes related to ROS. This excess in UPE may explain some phenomena of phosphene and visual hallucinations. However, due to the lack of available information about phosphene and visual hallucinations and the function of UPE in the visual cortex, more research and experiments are needed to investigate these phenomena and the factors affecting the production of UPE.

## Author Contributions

- Conceptualization: TE, AL, IB, VS
- Data curation:TE, AL, NK
- Formal analysis: TE, AL, NK
- Funding acquisition: MA, AS, CS, VS, DO
- Methodology: AL, MA, MN, NK, AS, TE
- Project administration: MN, TE, NK
- Resources: TE, NK, MA, MN, AS
- Supervision: TE, NK, VS
- Discussion: IB, AL, NK, HZH, CS, VS, DO
- Hypothesis: HZH, CS, IB, VS
- Writing – original draft: TE, AL, NK, IB, VS
- Writing – review and editing: TE, NK, CS, HZH, VS, DO

## Acknowledgments

We would like to thank the Histomorphometry and Stereology Research Center of Shiraz university of medical sciences for their Cooperation for doing this project. We also thank the Natural Sciences and Engineering Research Council (NSERC) of Canada and the National Research Council (NRC) of Canada.

## Data Availability Statement

The datasets are available based on request.

## References

1. Cifra, M. Pospisil, P. Ultra-weak photon emission from biological samples: definition, mechanisms, properties, detection and applications. J. Photochem. Photobiol. B. Biol 139, 2–10 (2014).

2. Pospisil, P., Prasad, A. Rac, M. Role of reactive oxygen species in ultra-weak photon emission in biological systems. Journal of Photochemistry and Photobiology B: Biology 139, 11–23 (2014).

3. Esmaeilpour, T., Fereydouni, E., Dehghani, F., Bókkon, I., Panjehshahin, M. R., Császár-Nagy, N., et al. An experimental investigation of ultraweak photon emission from adult murine neural stem cells. Sci. Rep. (2020) 10, 463

4. Salari, V., Scholkmann, F., Vimal, L. P., Csaszár, N., Aslani, M., and Bókkon, I. Phosphenes, retinal discrete dark noise, negative aferimages and retinogeniculate projections: a new explanatory framework based on endogenous ocular luminescence. Prog. Ret. Eye Res. (2017) 60, 101–119.

5. Salari, V., Valian, H., Bassereh, H., Bokkon, I., and Barkhordari, A. Ultraweak photon emission in the brain. J. Integ. Neurosci. (2015) 14, 419–429.

6. Isojima, Y., Isoshima, T., Nagai, K., Kikuchi, K. Nakagawa, H. Ultraweak biochemiluminescence detected from rat hippocampal slices. Neuroreport 6, 658–660 (1995).

7. Kobayashi, M. et al. In vivo imaging of spontaneous ultraweak photon emission from a rat’s brain correlated with cerebral energy metabolism and oxidative stress. Neurosci. Res. 34, 103–113 (1999).

8. Salari, V., Bókkon, I., Ghobadi, R., Scholkmann, F. Tuszynski, J. A. Relationship between intelligence and spectral characteristics of brain biophoton emission: Correlation does not automatically imply causation. P. Nat. Acad. Sci. 113(38), E5540–E5541 (2016).

9. Moszczynska A. Neurobiology and Clinical Manifestations of Methamphetamine Neurotoxicity. Psychiatr Times. 2016 Sep;33(9):16–18.

10. Courtney KE, Ray LA. Methamphetamine: an update on epidemiology, pharmacology, clinical phenomenology, and treatment literature. Drug Alcohol Depend. 2014 Oct 1;143:11–21.

11. Cruickshank CC, Dyer KR. A review of the clinical pharmacology of methamphetamine. Addiction. 2009; 104: 1085–1099.

12. Teodorof-Diedrich C, Spector SA. Human Immunodeficiency Virus Type 1 and Methamphetamine-Mediated Mitochondrial Damage and Neuronal Degeneration in Human Neurons. J Virol. 2020 Sep 29;94(20):e00924–20.

13. Cadet J. L., Jayanthi S., Deng X. Methamphetamine-induced neuronal apoptosis involves the activation of multiple death pathways. Review. Neurotoxicity Research, 8, 3–4, 199-206, 2005.

14. Yu S, Zhu L, Shen Q, Bai X, Di X. Recent advances in methamphetamine neurotoxicity mechanisms and its molecular pathophysiology. Behav Neurol. 2015;2015:103969.

15. Davidson C, Gow AJ, Lee TH, Ellinwood EH. Methamphetamine neurotoxicity: necrotic and apoptotic mechanisms and relevance to human abuse and treatment. Brain Res Brain Res Rev. 2001 Aug;36(1):1–22.

16. Berman S, O’Neill J, Fears S, Bartzokis G, London ED. Abuse of amphetamines and structural abnormalities in the brain. Ann N Y Acad Sci. 2008;1141:195–220

17. Graves SM, Schwarzschild SE, Tai RA, Chen Y, Surmeier DJ. Mitochondrial oxidant stress mediates methamphetamine neurotoxicity in substantia nigra dopaminergic neurons. Neurobiol Dis. 2021 Aug;156:105409.

18. Krasnova IN, Cadet JL. Methamphetamine toxicity and messengers of death. Brain Res Rev. 2009 May;60(2):379–407.

19. Sheikhi Koohsar J, Faeghi F, Rafaiee R, Niroumand Sarvandani M, Masjoodi S, Kalalian Moghaddam H. Metabolite Alternations in the Dopamine Circuit Associated with Methamphetamine-Related Psychotic Symptoms: A Proton Magnetic Resonance Spectroscopy Study. Iran J Psychiatry. 2022 Jan;17(1):91–98.

20. Wearne TA, Cornish JL. A Comparison of Methamphetamine-Induced Psychosis and Schizophrenia: A Review of Positive, Negative, and Cognitive Symptomatology. Front Psychiatry. 2018 Oct 10;9:491

21. Zarrabi H, Khalkhali M, Hamidi A, Ahmadi R, Zavarmousavi M. Clinical features, course and treatment of methamphetamine-induced psychosis in psychiatric inpatients. BMC Psychiatry. 2016 Feb 25;16:44.

22. Shariat SV, Elahi A. Symptoms and course of psychosis after methamphetamine abuse: one-year follow-up of a case. Prim Care Companion J Clin Psychiatry. 2010;12(5):PCC.10l00959.

23. Yui K, Goto K, Ikemoto S, Ishiguro T. Methamphetamine psychosis: spontaneous recurrence of paranoid-hallucinatory states and monoamine neurotransmitter function. J Clin Psychopharmacol. 1997 Feb;17(1):34–43.

24. Lamyai W, Pono K, Indrakamhaeng D, Saengsin A, Songhong N, Khuwuthyakorn P, Sribanditmongkol P, Junkuy A, Srisurapanont M. Risks of psychosis in methamphetamine users: cross-sectional study in Thailand. BMJ Open. 2019 Oct 14;9(10):e032711.

25. Juan CA, Pérez de la Lastra JM, Plou FJ, Pérez-Lebeña E. The Chemistry of Reactive Oxygen Species (ROS) Revisited: Outlining Their Role in Biological Macromolecules (DNA, Lipids and Proteins) and Induced Pathologies. International Journal of Molecular Sciences. 2021;22(9):4642. PubMed PMID: doi:10.3390/ijms22094642.

26. Tsikas D. Assessment of lipid peroxidation by measuring malondialdehyde (MDA) and relatives in biological samples: Analytical and biological challenges. Analytical biochemistry. 2017 May 1;524:13–30. PubMed PMID: 27789233. Epub 2016/10/30. eng.

27. Gawel-S, Wardas M, Niedworok E, Wardas P. [Malondialdehyde (MDA) as a lipid peroxidation marker]. Wiadomosci lekarskie (Warsaw, Poland : 1960). 2004;57(9-10):453–5. PubMed PMID: 15765761. Epub 2005/03/16. Dialdehyd malonowy (MDA) jako wskáznik procesów peroksydacji lipidów w organizmie. pol.

28. Ayala A, Muñoz MF, Argüelles S. Lipid peroxidation: production, metabolism, and signaling mechanisms of malondialdehyde and 4-hydroxy-2-nonenal. Oxidative medicine and cellular longevity. 2014;2014:360438. PubMed PMID: 24999379. Pubmed Central PMCID: PMC4066722. Epub 2014/07/08. eng

29. Jesus Aguilar Diaz De Leon, Chad R Borges, Evaluation of Oxidative Stress in Biological Samples Using the Thiobarbituric Acid Reactive Substances Assay. J Vis Exp 12(159), 2020.

30. Cristina Mas-Bargues, Consuelo Escriva, Mar Dromant, Consuelo Borras, Jose Vina. Lipid peroxidation as measured by chromatographic determination of malondialdehyde. Human plasma reference values in health and disease. Archives of Biochemistry and Biophysics, 709 (2021) 108941.

31. Graham DG. Oxidative pathways for catecholamines in the genesis of neuromelanin and cytotoxic quinones. Mol Pharmacol. 1978;14(4):633–43.

32. hassani Moghaddam M, Boroujeni ME, Vakili K, Fathi M, Abdollahifar M-A, Eskandari N, et al. Functional and structural alternations in the choroid plexus upon Methamphetamine exposure. Neuroscience Letters. 2021:136246.

33. Fleckenstein AE, Wilkins DG, Gibb JW, Hanson GR. Interaction Between Hyperthermia and Oxygen Radical Formation in the 5-Hydroxytryptaminergic Response to a Single Methamphetamine Administration. Journal of Pharmacology and Experimental Therapeutics. 1997;283(1):281.

34. Nakato Y, Abekawa T, Ito K, Inoue T, Koyama T. Lamotrigine blocks apoptosis induced by repeated administration of high-dose methamphetamine in the medial prefrontal cortex of rats. Neuroscience letters. 2011;490(3):161–4.

35. Cadet JL, Jayanthi S, Deng X. Speed kills: cellular and molecular bases of methamphetamine[U+2010]induced nerve terminal degeneration and neuronal apoptosis. The FASEB Journal. 2003;17(13):1775–88.

36. Kuhn DM, Francescutti-Verbeem DM, Thomas DM. Dopamine Quinones Activate Microglia and Induce a Neurotoxic Gene Expression Profile. Annals of the New York Academy of Sciences. 2006;1074(1):31–41.

37. Bókkon I, Salari V, Tuszynski JA, Antal I. Estimation of the number of UPEs involved in the visual perception of a single-object image: UPE intensity can be considerably higher inside cells than outside. Journal of Photochemistry and Photobiology B: Biology. 2010;100(3):160–6.

38. Young N, Collins C, Kaas J. Cell and neuron densities in the primary motor cortex of primates. Frontiers in Neural Circuits. 2013;7(30).

39. Stefanatos R, Sanz A. The role of mitochondrial ROS in the aging brain. FEBS letters. 2018;592(5):743–58.

40. Boroojerdi B, Meister IG, Foltys H, Sparing R, Cohen LG, Töpper R. Visual and motor cortex excitability: a transcranial magnetic stimulation study. Clinical Neurophysiology. 2002;113(9):1501–4.

41. Kähkönen S, Wilenius J, Komssi S, Ilmoniemi RJ. Distinct differences in cortical reactivity of motor and prefrontal cortices to magnetic stimulation. Clin Neurophysiol. 2004;115(3):583–8.

42. Caulfield KA, Li X, George MS. A reexamination of motor and prefrontal TMS in tobacco use disorder: Time for personalized dosing based on electric field modeling? Clin Neurophysiol. 2021;132(9):2199–207.

43. Isojima Y, Isoshima T, Nagai K, Kikuchi K, Nakagawa H. Explanations step by step about Bókkon’s biophysical picture representation model (also called intrinsic biophysical virtual visual reality) during visual perception and imagery. Methods.93:163–8.

44. Fassbender C, Lesh TA, Ursu S, Salo R. Reaction time variability and related brain activity in methamphetamine psychosis. Biol Psychiatry. 2015 Mar 1;77(5):465–74.

45. Dennis K. Miller et al. “Subchronic apocynin treatment attenuates methamphetamine-induced dopamine release and hyperactivity in rats”. Life Sciences 98.1 (2014), 6–11.

46. Hadi Zadeh-Haghighi and Christoph Simon. “Entangled radicals may explain lithium effects on hyperactivity”. Scientific Reports 11.1 (2021).

47. Helmut Sies and Dean P. Jones. “Reactive oxygen species (ROS) as pleiotropic physiological signalling agents”. Nature Reviews Molecular Cell Biology 21.7 (2020), 363–383.

48. P. J. Hore and Henrik Mouritsen. “The Radical-Pair Mechanism of Magnetoreception”. Annual Review of Biophysics 45.1 (2016), 299–344.

49. Tianzhen Chen et al. “A transcranial magnetic stimulation protocol for decreasing the craving of methamphetamine-dependent patients”. STAR Protocols 2.4 (2021), 100944.

50. Qingming Liu et al. “Either at left or right, both high and low frequency rTMS of dorsolateral prefrontal cortex decreases cue induced craving for methamphetamine”. The American Journal on Addictions 26.8 (2017), 776–779.

51. Qiongdan Liang et al. “Intervention Effect of Repetitive TMS on Behavioral Adjustment After Error Commission in Long-Term Methamphetamine Addicts: Evidence From a Two-Choice Oddball Task”. Neuroscience Bulletin 34.3 (2018), 449–456.

52. Tianye Ma, Yurong Sun, and Yixuan Ku. “Effects of Non-invasive Brain Stimulation on Stimulant Craving in Users of Cocaine, Amphetamine, or Methamphetamine: A Systematic Review and Meta-Analysis”. Frontiers in Neuroscience 13 (2019).

53. Jiajin Yuan et al. “Effect of Low-Frequency Repetitive Transcranial Magnetic Stimulation on Impulse Inhibition in Abstinent Patients With Methamphetamine Addiction”. JAMA Network Open 3.3 (2020), e200910.

54. Hadi Zadeh-Haghighi and Christoph Simon. “Magnetic field effects in biology from the perspective of the radical pair mechanism”. Journal of The Royal Society Interface 19.193 (2022).

